# Smoothing over “rough” mismanagement: establishing protective harvest limits for native nongame fishes

**DOI:** 10.64898/2026.02.24.707793

**Authors:** Alexandria N. Ginez, Solomon R. David, Alec R. Lackmann, Bonnie J.E. Myers, Tyler J. Winter, Robert A. Lusardi, Andrew L. Rypel

## Abstract

There is growing interest in establishing more protective regulations for native fishes historically classified as “rough fish”- a term ascribed to species of low-to-zero perceived commercial value. Yet high-quality population data are lacking for most species and populations, precluding determination of sustainable harvest limits using standard methods. Here, we present an inductive and ecosystem -based approach for comparing and aligning harvest limits of diverse fish species. Our approach centers on the production/biomass (P/B) ratio as the key instrument for gauging sustainable harvest. P/B is the biomass turnover rate in populations and therefore quantifies the return rate of any removed biomass in populations. We extracted and summarized data from existing studies, representing a total of 517 empirical estimates of secondary production, biomass, and P/B ratios. We subsequently developed a highly predictive statistical model (R^2^ = 0.90), demonstrating P/B is largely a function of maximum age across species. We then developed a separate database on age, growth, and longevity data for most native fishes of interest across the USA. For each species and population, we leveraged the above statistical model to predict and compare mean P/B across species. Results show most native fishes express P/B values similar to, or lower than, traditional game fish species. Accordingly, harvest limits across species groups can be harmonized with those of other managed species. For example, native nongame species like Bigmouth Buffalo *Ictiobus cyprinellus* and Freshwater Drum *Aplodinotus grunniens* are long-lived with slow replacement rates that are statistically clustered with those observed in Lake Sturgeon *Acipenser fulvescens* and trophy Muskellunge *Esox masquinongy* populations, two popular game fish species. Harvest limits for these nongame species would therefore need to be similarly low for these species to ensure comparable sustainability. To understand broad patterns of harvest limit alignment, we modeled relationships between daily bag limits of managed species and P/B for five test states. Model uniformly showed non-linear trends with high residuals (suggesting excessive bag limits) common for panfish species and low residuals (suggesting overly conservative bag limits) common for trout species. Managers can use the results of this study to estimate harvest limits native fishes.

## Introduction

Freshwater ecosystems are degrading at unprecedented rates (Carpenter et al. 2011; Sayer et al. 2025), and the status of native fishes exemplifies these trends (Warren and Burr 1994; Albert et al. 2021; Leidy and Moyle 2021). Only about half of Earth’s freshwater fish species are considered stable; the remainder are either declining toward extinction or considered data deficient (Sayer et al. 2025). However, in many regions, the status of native fishes and the severity of their collapse is alarming (Contreras-Balderas et al. 2008; Moyle et al. 2011). There are myriad factors driving fish population declines (Duncan and Lockwood 2001; Arthington et al. 2016); however, mismanagement of natural resources, including allowing unlimited harvest and concomitant overfishing leading to population loss remains a major issue (Post et al. 2002; Allan et al. 2005; Fluet-Chouinard et al. 2018; Embke et al. 2019).

There is rising awareness that historical fisheries management systems have largely failed in protecting most native fishes (Guy et al. 2021; Rypel et al. 2021; Clancy et al. 2025). “Rough fish” (hereafter referred to as “native nongame species” unless referencing historical usage) is a term given to a large group of native North American fish species. These are fishes perceived to have low-to-zero recreational and commercial value and therefore receive little-to-no conservation management actions (Scarnecchia 1992; Kelly and David 2024; Winter 2024). Previous studies provide comprehensive reviews on the history of inland fisheries management (Nielson 1999), including how native nongame management specifically has languished (Montague et al. 2023; Lackmann et al. 2024a). In some cases, management has been actively antagonistic. For example, it was once common practice to poison waterbodies to remove all native fishes to restock with non-native game fishes (Rose and Moen 1953; Sandoz 1960; Hubbs 1963; Binns 1965). Such management actions are largely unheard of today; however, vestiges of these past perspectives remain in the form of unlimited harvest restrictions for many species, lack of wanton waste laws, and miniscule data collection (Lackmann et al. 2019; Scarnecchia and Schooley 2020; Lackmann et al. 2024). Indeed, there is now recognition that native nongame species also often benefit from management (Smith et al. 2020; Jacinto et al. 2023), and that some species hold value independent of traditional recreational angling (Cooke et al. 2020; Reid et al. 2021). For example, Indigenous tribes have always valued suckers (family Catostomidae), and these fishes can play a central role in tribal cultures (Royle 2021; Rypel et al. 2021; Shelley and Nicola 2022). Furthermore, perspectives change over time. Some species previously classified as “rough fish” (e.g., catfishes [Ictaluridae], Alligator Gar *Atractosteus spatula*, sturgeon [Acipenseridae]) developed devout angling communities over the last 10-50 years. In most cases, these species are now considered game species and receive adequate management attention (Pikitch et al. 2005; Porath et al. 2021; Rypel et al. 2021). Yet many more native fishes remain largely unregulated (e.g., unlimited bag limits) and receive minimal or no management focus, despite growing exploitative fisheries on many of the taxa including buffalofishes (*Ictiobus* spp.; Lackmann et al. 2019, 2021, 2024c), carpsuckers (*Carpiodes* spp.; Lackmann et al. 2022a, in press), bowfin (*Amia* spp.; Sinopoli and Stewart 2021; Lackmann et al. 2022b), redhorses (*Moxostoma* spp.; Lackmann et al. 2024b), Freshwater Drum *Aplodinotus grunniens* (Neely et al. 2024), and others.

There are real obstacles to improved conservation management of native fishes in the United States. These include: (1) traditional funding models that emphasize game species (Clarkson et al. 2005; Scarnecchia et al. 2021); and (2) lack of information on the ecology of native fishes to inform improved management (Clark and May 2002; Guy et al. 2021). In terms of funding, nearly every US state acquires funds for fisheries management activities through a combination of fishing license sales and Sport Fish Restoration (SFR) or Dingell-Johnson Act funds-a federal excise tax on fishing items, boats and boat gas (Scarnecchia et al. 2021). For SFR, funds flow back to the states via the US Fish and Wildlife Service, but only in proportion to their fishing license sale totals. Therefore, funds to support fisheries management flow directly from anglers and the fishing industry. This funding model has been an engine for strong and effective fisheries management in the US for decades (Börk 2024; Kopaska 2025), but only for the species for which management and the fishing industry focus (Scarnecchia et al. 2021). Therefore, one critique of this conservation model is that it does not advance management of species outside recreational angling value. Even for species of high interest to smaller and devoted groups of anglers, management activities for popular gamefishes are usually prioritized (Cooke et al. 2020; Clancy et al. 2025).

Most native nongame species historically received no funding, and subsequently our understanding of their ecology remains non-existent or incomplete at best (Donaldson et al. 2011; Guy et al. 2021). These knowledge gaps, in turn, preclude strong science-based management for native fishes and their habitats (Sass et al. 2017; Williams et al. 2020), and may exacerbate pre-existing misconceptions (Scarnecchia 1992). It is helpful to recognize that this problem is not novel. Marine fisheries are similarly complex and managed across huge species complexes expressing divergent and poorly understood life-histories (May et al. 1979; Murawski 1991; Kindsvater et al. 2016). These challenges led to creative management practices in the absence of detailed species and population information (e.g., Geromont and Butterworth 2015; Bridges et al. 2023). Sometimes termed ‘Robinhood’ or ‘data-limited’ approaches, these methods use available data from some species to inform assessments of others (Punt et al. 2011; Dowling et al. 2015). The key innovation is that such approaches support timely decision-making, rather than waiting until populations decline further. Though widely applied in marine fisheries management, similar approaches are uncommon in freshwater fisheries (but see Tracy et al. 2025).

Ecosystem ecology provides an overlooked theoretical foundation through which sustainable harvest can be better understood. For example, fish secondary production is defined as the rate at which heterotrophic biomass is generated (Waters 1977; Benke 2010). If harvest rates for any fishery exceed production, biomass will decline (Waters 1992; Rypel et al. 2015). And because production and biomass are interrelated, the ratio of the two describes the quotient of production independent of biomass. The production/biomass or P/B ratio is therefore mathematically the biomass replacement rate (Banse and Mosher 1980; Stites and Benke 1989; Layman and Rypel 2020). Measured in units of inverse time, P/B provides a precise measure of the return rate for biomass in any population. For example, P/B ≈ 1.00 (e.g. common with trout populations) indicates that biomass turns over approximately annually, whereas P/B ≈ 0.20 (common with Walleye *Sander vitreus*) indicates a population where biomass turns over roughly once every five years (Rypel et al. 2015; Myers et al. 2017; Rypel and David 2017). Populations with high P/B can generally tolerate higher levels of harvest (assuming sufficient biomass) whereas those with low P/B are more vulnerable to fishing (Rypel et al. 2015; Myers et al. 2017; Rypel and David 2017). When B is low, a collapse can occur especially fast, but when B is high, declines or collapse can be slower, as B is successively worked down (Embke et al. 2019). Blue whales *Balaenoptera musculus* have a P/B = 0.02 (Christensen and Walters 2024), meaning their biomass turns over just once every 50 years. Virtually any harvest mortality on top of natural mortality for a species like this likely invites biomass to decline and, in this case, helps contextualize historical whale declines (McHugh 1977). Though rarely acknowledged, most sustainable harvest models in fisheries derive from these metrics. For example, the first chapter of Walters and Martell (2004) is devoted to describing these conceptual connections and providing the relevant derivatives to modern stock assessment tools. Initial tests of proportional stock density (PSD) metrics were essentially patterned after empirical production measures taken by Swingle (1950) in experimental ponds (Willis et al. 1993). P/B also remains a central and prominent parameter in the build and deployment of ecosystem-based food web models like Ecopath and Ecosim (Christensen and Walters 2004; Heymans et al. 2016).

Unfortunately, empirical studies of fish production are comparatively rare, likely because of the effort and cost needed to acquire all the component pieces for its direct estimation (Rypel and David 2017). Most biologists elect to study the individual components themselves, e.g., population numbers, growth, or possibly biomass (Rypel et al. 2015). However, there are sufficient studies of production dynamics in game and some nongame fish populations to understand the broader drivers of production. For example, what are the population and life-history drivers of P/B in fishes? If a general relationship could be uncovered, there is in turn strong potential to apply these patterns to better understand the conservation needs of many native fishes, even with limited data.

The goals of this study are to (1) understand the relationship between P/B and longevity across freshwater fishes; (2) utilize the statistical relationship developed above to predict P/B for all species classified as “rough fish” in Rypel et. al. (2021); and (3) model management alignment of game fishes and native nongame fishes in a series of test states (GA, OK, KY, NY, MN). Ultimately, we present this study as one potential method for advancing conservation and management of a suite of native nongame fishes in the USA, specifically through the development of more protective harvest regulations.

## Methods

### Meta-analysis

We first conducted a meta-analysis of secondary production studies in freshwater fishes to evaluate how select fish traits shape P/B. The meta-analysis focused on studies that contained sufficient information on secondary production, biomass, P/B, size- and age-structure of fish populations. Secondary production data are available for most species traditionally classified as game fish; many of these species have replicated estimates of production across ecosystems and years (Supplementary Dataset 1). However, there is also good availability of secondary production data for some native nongame species, especially stream fishes (e.g., Myers et al. 2017). Methods of secondary production estimation varied across studies, and included the instantaneous growth method, the size-frequency method and the increment summation method. We viewed all these methods as equally legitimate estimates of production; however, we excluded production estimates derived from empirical models. In some cases, we developed new estimates of production, biomass and P/B from published studies using methods outlined in Rypel et al. (2018). In these cases, the variables required to calculate production were collected but not analyzed, since it was not the study’s objective.

We developed a statistical relationship between P/B and maximum longevity of fishes using weighted log-linear regression. A mean P/B value was calculated for each species using the empirical meta-analysis described above. These values were then plotted against mean longevity (maximum age) for each species from the same studies used to estimate P/B, and a weighted log-linear regression fit to these values. In the model, the dependent variable was log_10_(P/B), log_10_(maximum age) was the independent variable, and sample size of computed means was the weighting factor. Thus, our regression gave more weight to species with higher numbers of independent P/B estimates. Analogous models were built to test the relationships between 1) P/B and mean age; 2) P/B and average maximum size (Western 1983) (assayed as L_inf_ from the Von Bertalanffy Growth Function); and 3) P/B and growth (assayed as the k parameter from the Von Bertalanffy Growth Function). We also explored different data filtering methods for studies used in our calculations including 1) using species data only if the values were represented by more than one unique study (hereafter “species with replication”) or 2) utilizing all available data. From these life-history characters and data selection methods, and because of the strength of this relationship, we ultimately focused exclusively on the P/B-to-longevity relationship for species with replication. This decision is further supported because: a) it aligns with strong relationships documented in prior studies (Robertson 1979; Kwak and Waters 1997; Randall and Minns 2000); b) it is an uncomplicated regression model that can be used by any manager; and c) maximum age is a simple parameter to collect and can be routinely generated during fisheries surveys (Casselman et al. 1999; Rypel 2012; Lackmann et al. 2019).

We then conducted a second meta-analysis focused on growth and longevity. Age and growth data were collated for both game species and native nongame species historically considered “rough fish” as identified in Rypel et al. (2021). Longevity data were collected from studies across the geographic range of each species, and only from studies that utilized validated age-estimation structures such as otoliths, vertebrae, fin rays, (Beamish and McFarlane 1983). Age data from scales were not accepted for any species due to inconsistencies in age estimation (Goeman et al. 1984; Maceina et al. 2007; Quist et al. 2023). Once completely assembled, we predicted P/B based on maximum age for all game and native nongame fishes lacking empirical values using the weighted log-linear statistical model described above. Predicted P/B data from maximum ages were then combined with empirically derived P/B data to develop one unified database of P/B values across species (Supplementary Dataset 2). We then calculated and compared mean P/B values across all species. A k-means cluster analysis was then used, using the mean P/B values for each species of interest. This cluster analysis provided four groups of fishes with relatively similar biomass replacement rates, which we interpret as those that would benefit from similar harvest limits.

Finally, we explored the extent to which mean P/B values for game fish species explain current daily bag limits for five select states: MN, GA, OK, KY, NY, and all states combined. These states were chosen to represent a cross-section of latitudes and fisheries encompassing the geographic ranges of many of our study species. These analyses are therefore not intended to be comprehensive, or even suggestive of what each of these states should do. Rather, we present them as examples of alignment exercises that can be used as a first step toward improved protections for native fishes based on current management practices. In each study state, we developed a weighted generalized additive model (GAM) that predicted bag limits for fishes in each region based on P/B. We selected the GAM modeling framework because it is agnostic to the shape of the statistical relationship and can accommodate a wide range of nonlinear patterns (Wood 2017). Therefore, in each state model, the state’s bag limits for each species represented the dependent variable, P/B for those same species was the explanatory variable, and P/B sample size was again a weighting factor. “Bag limits” discussed in this model and throughout the paper refer to the daily creel limit, or number of fish legally harvestable in a calendar day. Bag limit values were extracted from statewide fishing regulations posted in 2025 (Supplementary Data 3). The models do not consider any other methods of harvest regulations or restrictions such as quotas, size restrictions or slot limits, sex restrictions, seasonal closures, or gear restrictions as they require a more detailed understanding of a population’s dynamics than longevity values (Gresswell and Harding 1997; Lennox et al. 2016; Haase et al. 2022). Longevity values used to compute estimated P/B values in each state exercise were constrained to those collected from only the region of interest or nearby using a k-means clustering on median latitude values of each longevity value’s study site. Residuals of all six GAMs were extracted and visualized across species to observe any patterns in current regulations and productivity-informed expectations for alignment considerations.

## Results

Our study incorporated data from 301 studies representing 108 freshwater fish species and 1,364 populations across the United States. A total of 32 species evaluated in this study are managed as game fish species, 37 species are native nongame fish, and the remaining 10 species are considered “rough fish escapees”, i.e., species once considered “rough fish” that are now managed as game species such as Alligator Gar and Lake Sturgeon *Acipenser fulvescens* (Rypel et al. 2021).

For our secondary production meta-analysis, we collated a total of 470 P/B values from 48 species representing 516 populations. Across all species, empirical P/B ranged 0.03 (or once every 39 years, Lake Sturgeon) to an incredible 18.16 (or 18 times a year, Amargosa Pupfish *Cyprinodon nevadensis*); however mean P/B (calculated as the mean of means) across all species was 0.62 ± 0.94 SE. Yet P/B also varies considerably within species. For example, across 218 Walleye populations, P/B ranged 0.04-0.65, and across 58 Brook Trout *Salvelinus fontinalis* populations, P/B ranged 0.41-2.79.

There was a significant relationship between log_10_(P/B) and log_10_(maximum age) when observing both species with replication (Figure 1f, weighted regression, log(P/B) = 1.68+(−1.14*(log(maximum age)), F = 110.1, R^2^ = 0.90, P < 0.0001) and all available species (Figure 1c, weighted regression, log (P/B) = 0.92+(−0.83*(log(maximum age)), F = 115.5, R^2^ = 0.71, P < 0.0001). The relationship between log_10_(P/B) and log_10_(L_inf_) was weaker (Figure 1b, weighted regression, log(P/B) = 5.03+(−0.96*(log(L_inf_)), F = 15.7, R^2^ = 0.48, P = 0.0013 and Figure 1e, weighted regression, log(P/B) = 7.56+(−1.36*(log(L_inf_)), F = 9.37, R^2^ = 0.48, P = 0.0156). For both data selection methods, there was also a significant relationship between K and P/B (Figure 1a, weighted regression, log(P/B) = 1.08+(1.23*(log(K)), F = 25.32, R^2^ = 0.60, P = 0.0001 and Figure 1d, weighted regression, log(P/B) = 1.7+(1.60*(log(K)), F = 0.63, R^2^ = 0.63, P = 0.0036).

**Figure 1.**
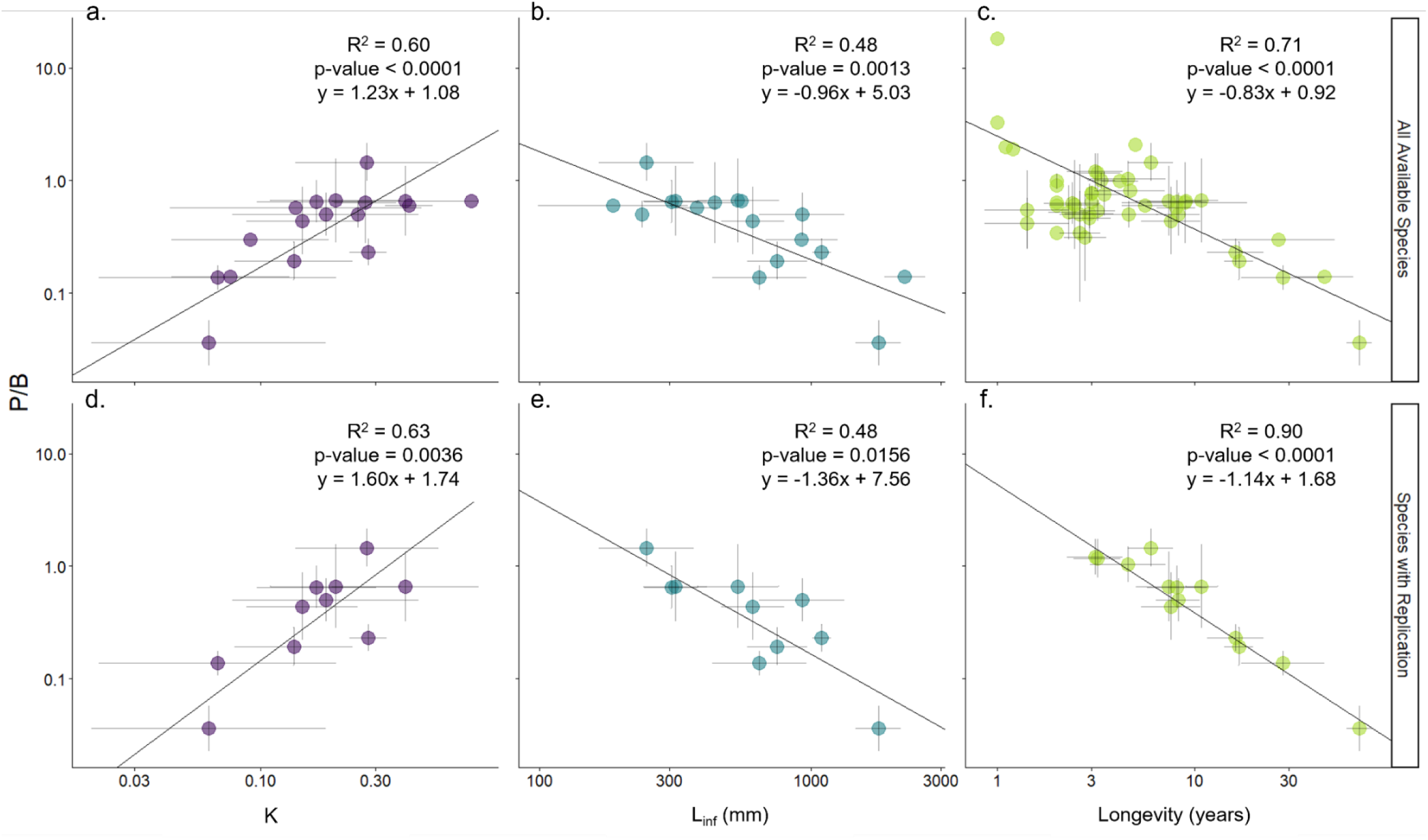
Weighted linear regressions highlighting statistical relationships between P/B and K (1a., 1d.), Linf (1b., 1e.), and P/B and Longevity (1c., 1f.). Note that relationships are repeated using two different data selection and filtering methods. Each point represents an average for each species ± 1 SE.

In our meta-analysis of fish age structure and growth rates, we collated information on 75 species, representing a total of 851 fish populations. Maximum longevity for the target species ranged 2 (Black Bullhead *Ameirus melas*) to 148 y (Bigmouth Buffalo *Ictiobus cyprinellus*), with mean maximum age = 20 y ± 1.02 SE. When the weighted regression described above was applied to these data, it produced a similarly wide range of estimated P/B values as those which were empirically-derived. For example, estimated P/B values based on maximum age ranged 0.02-5.37 with a mean P/B = 0.52 ± 0.01 SE (Figure 2).

**Figure 2.**
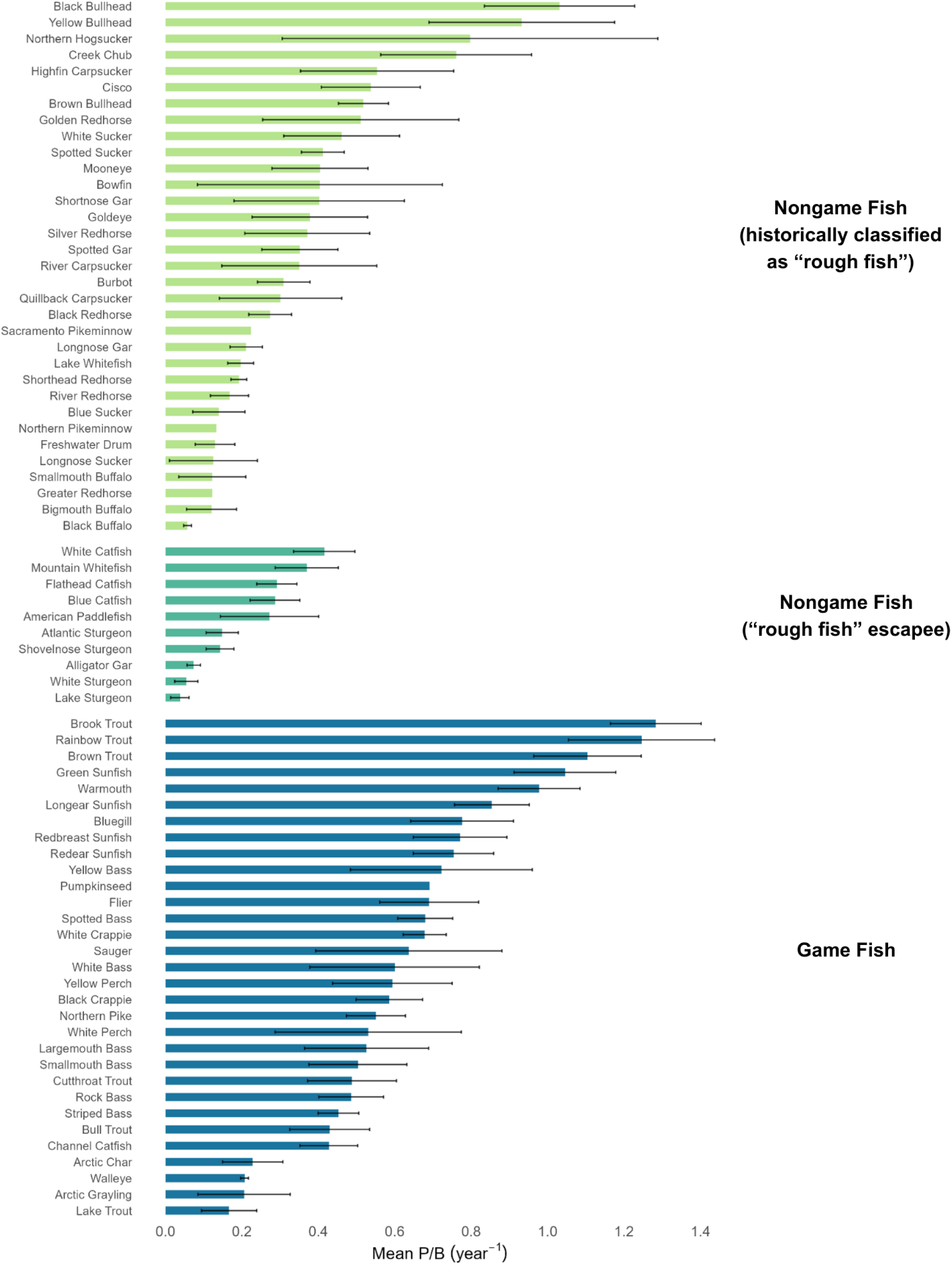
Mean P/B ± 1 SE for three groups of fishes presented in rank order by group. P/B values presented in this figure represent both empirical production estimates and modeled P/B estimates using the statistical relationship based on longevity present in Figure 1f (see Methods).

In general, there was a similar or lower range of P/B estimates for species managed as native nongame versus game fish (Figure 2). For example, redhorse (*Moxostoma spp.*) and Blue Sucker *Cycleptus elongatus* have estimated P/B values (0.13-0.20) similar to those observed for Walleye and Lake Trout *Salvelinus namaycush*. Species such as White Sucker *Catostomus commersonii* have similar values to game fish such as Rock Bass *Ambloplites rupestrus* and Channel Catfish *Ictalurus punctatus* (∼0.45). However, some native nongame species have extremely low P/B. Average P/B for Black Buffalo *Ictiobus niger* is 0.05, Bigmouth Buffalo at 0.10, with Smallmouth Buffalo *Ictiobus bubalus*, Longnose Sucker *Catostomus catostomus*, and Freshwater Drum at around 0.13. These mean P/Bs are lower than any game fish species examined in this study. Lake Sturgeon and White Sturgeon *Acipenser transmontanus* have the lowest P/B values of any species examined in this study (0.04-0.05), however they are considered “rough fish escapees” (Rypel et al. 2021).

Our weighted GAMs produced significant non-linear models for every state and region examined. Therefore, daily bag limits for most species in each region can be predicted based on P/B in a way that aligns with current regulations. At moderate P/Bs, bag limits generally plateau as a function of P/B in state-specific GAMs. However, the combined GAM representing a regional pattern depicts a marked increase in bag limits. At high P/Bs, bag limits were lower than expected if the linear trend at moderate P/Bs was continued. Residuals around the GAMs further contextualize some important patterns and case studies. For example, panfish species (Centrarchidae), almost uniformly have harvest limits of 10 fish per day or greater in most states, but these species also have high residuals in the GAMs. Thus, harvest limits for panfish are higher than predicted by the GAMs (Figure 3). In contrast, black bass (*Micropterus* sp.) and trout harvest limits show the opposite pattern; harvest limits are lower than predicted by GAMs. These species, in turn, help shape the consistent functional response curve shapes observed in the GAMs. The residuals demonstrate deviation between the implemented bag limits and productivity-informed expectations, so residual magnitude represent policy schema rather than literal differences in policy and productivity-based harvest.

**Figure 3.**
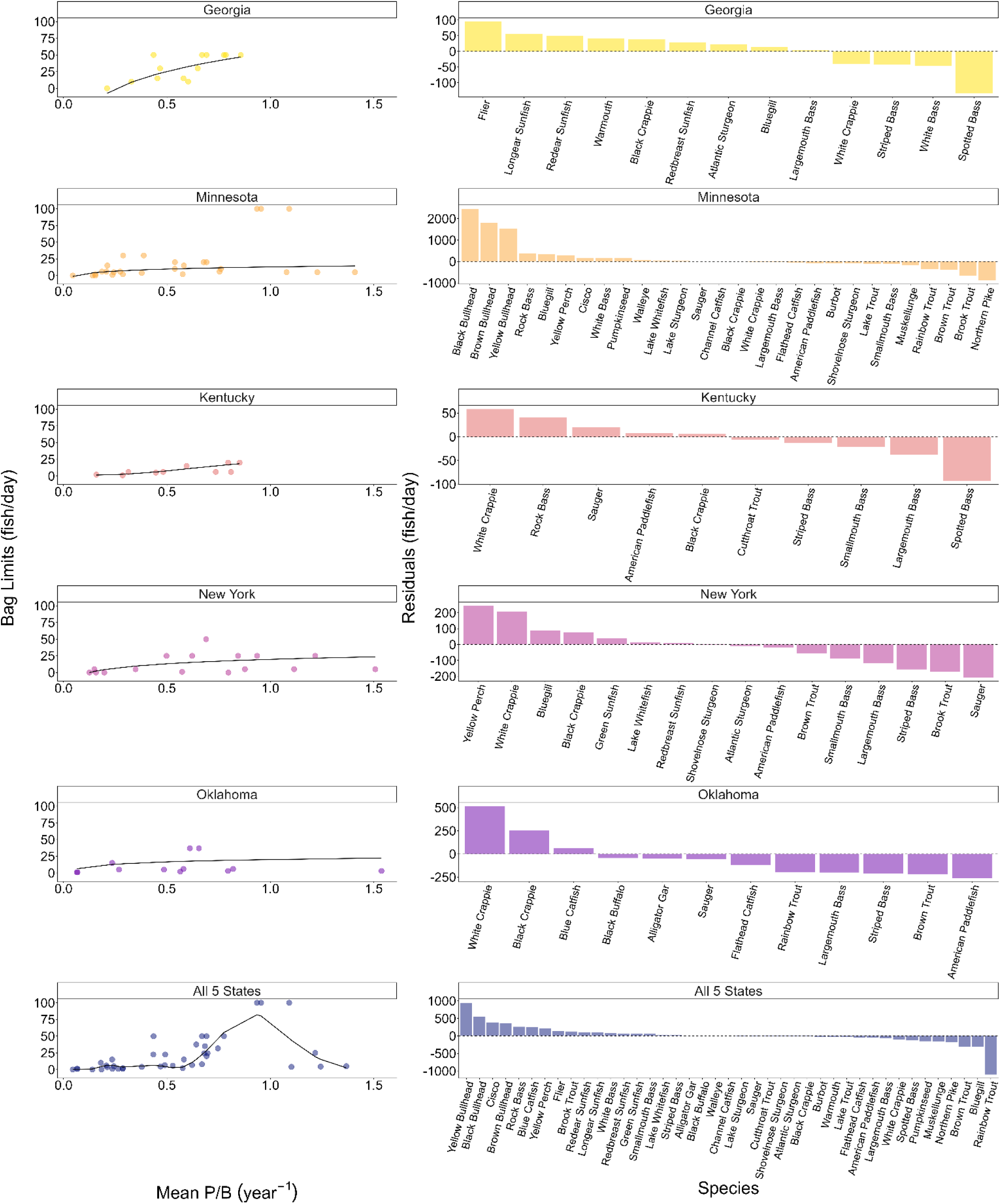
(Left) Weighted GAMs illustrating non-linear relationships between species P/B and bag limits for 5 select states (GA, MN, KY, NY, and OK) and all states combined. (Right) GAM residuals for each region, presented in rank order (highly positive residuals = higher bags than predicted by model, highly negative residuals = lower bags than predicted by model).

## Discussion

Native nongame fishes classified as “rough fish” are neglected but integral components of fisheries and ecosystems (Scarnecchia 1992; Rypel et al. 2021). As such, there are increasing calls for states to sustainably manage these and other species in more effective and respectful ways (Montague et al. 2023; Murchie et al. 2024; Winter 2024). One of the main hurdles to this goal, however, is the lack of information on life-history and species ecology sufficient to make critical management determinations (Guy et al. 2021; Lackmann et al. 2023). In this study, we provide an inductive method for aligning harvest limits of native fishes with those commonly used for managed sportfishes. The cornerstone to our approach is the estimation of biomass replacement rate (P/B), which we show is conceptually and fundamentally linked to maximum age in fish populations. This approach, or one that is similar, has potential to advance conservation management of diverse native fishes, especially in situations where data are limited.

Maximum age is a fundamental property of fish populations, and our meta-analysis reinforces robust literature on this topic (Bonner 1965; Waters 1977; Banse and Mosher 1980). In perhaps the simplest heuristic for this study, turnover rate for long-lived fishes (e.g., sturgeons) must be much slower than short-lived ones (e.g., pupfishes [Cyprinodontidae]). Indeed, in previous studies of secondary production rates, lifespan was frequently correlated with P/B in significant ways (Randall and Minns 2000; Kwak and Waters 1997). These critical differences are evident in simple production calculation tables comparing long- and short-lived species. At advanced ages, while still positive, there is very little accrual of biomass occurring for long-lived species during any given year, and often for long time intervals. Biomass can be added over long time periods, but annual fixation rates are very low, and occasionally even negative. Thus, removal of just one of these individuals requires considerable time to replace. These observations are supported by results from age-structured population models that often project low levels of fishing mortality to maintain long-lived species (e.g., often just 0-5%) (Francis 1992; TenBrink and Spencer 2013; Smith et al. 2018; Blackburn et al. 2019; Parker et al. 2025; Kopf et al. 2025). Furthermore, in at least one example, outputs from production -based approaches and population modeling generate virtually identical results (Tsehaye et al. 2016; Rypel et al. 2018).

This study provides important evidence linking fish population age structure together with patterns in P/B, and by extension, implications for sustainable fisheries management. Analogous studies in marine fisheries similarly show how life-history strategy, but especially longevity, correlates with sustainable fisheries management practices (Berkeley et al. 2004; Heppell et al. 2005; Kindsvater et al. 2016). While none of the marine studies directly address the link between longevity and P/B per se, these studies ultimately make related points. That is, that species and populations can be broadly categorized into groups based on life-history and inherent vulnerability to improve management. Further, these groups can be used to understand which species and stocks might benefit from certain kinds of management, namely differing degrees of harvest.

Simple comparisons of P/B among game fish and native nongame fish are a powerful tool for thinking about how to better align harvest limits. It is interesting and likely useful that native nongame species largely express a similar range in P/B compared to game species. Therefore, aligning harvest limits might be as simple as comparing species in an analogous manner as that in Figure 4. A caveat is that low P/B species are more represented within the native nongame fish group, especially for the buffalofishes, some other suckers, gars, and Freshwater Drum. Consequently, harvest limits for these species are likely to be lower. In our analysis, species with P/B values over 0.65 tend to be game fish (Figure 2, Figure 4). Bullheads (*Ameirus* spp.), Northern Hogsucker *Hypentelium nigricans*, and Cisco *Coregonus artedi* had P/B values most comparable to species like Brown Trout *Salmo trutta* and Rainbow Trout *Oncorhynchus mykiss* (≥1.0).

**Figure 4.**
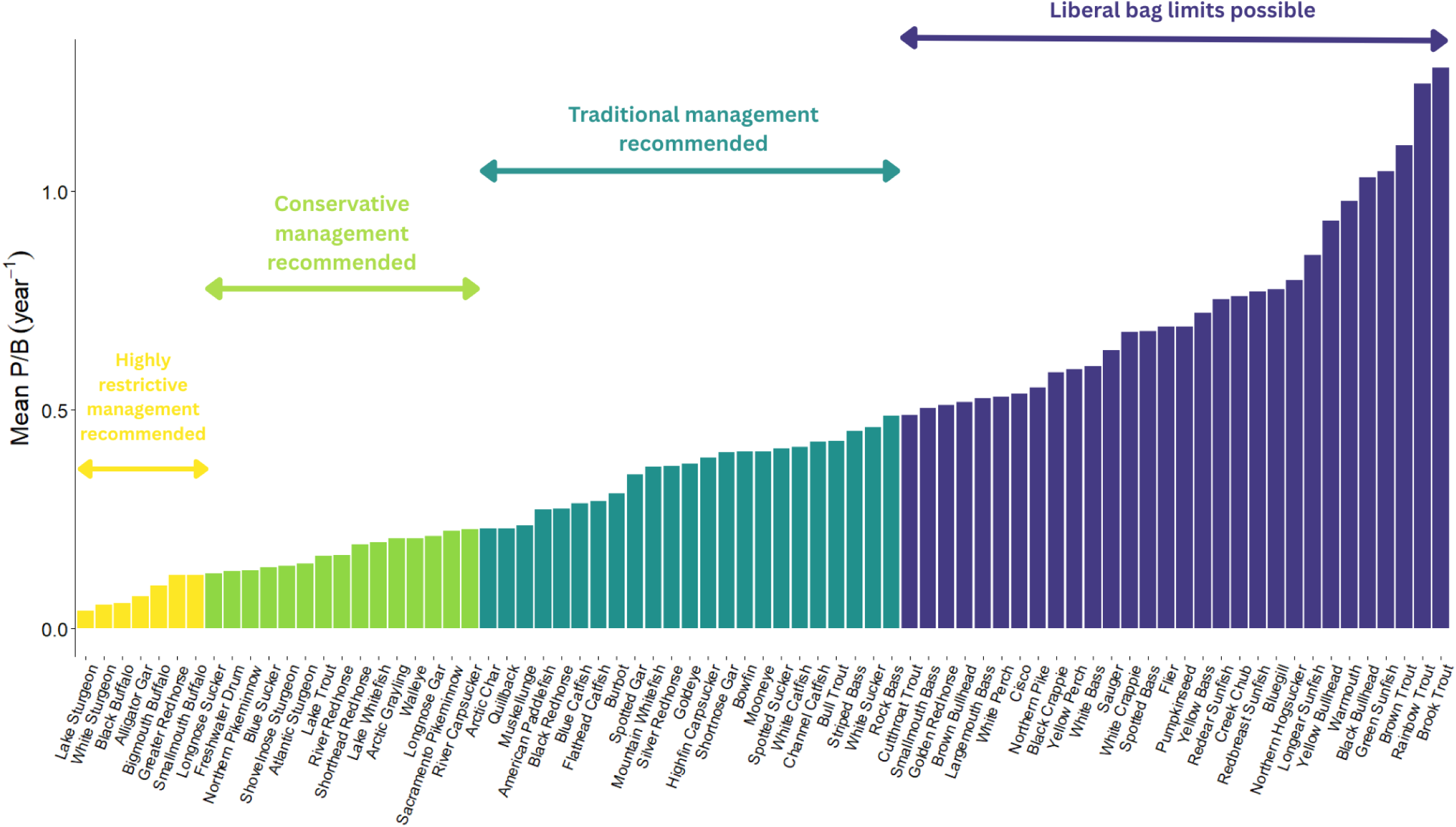
Results of cluster analysis on species P/B values. In this example, we label each cluster with potential management recommendations to clarify for managers where each species generally sits on the continuum.

One critique sometimes raised regarding reducing harvest limits on native fishes is that there is little direct evidence that fishing is having a direct effect. For example, in some locations there are seemingly plentiful numbers of buffalofishes and Freshwater Drum regularly caught by anglers. Yet as seen from the P/B values for these species, replacement rates for these species are clearly very low. This paradox is likely explained through high baseline standing stock biomass and the demographic storage effect (Winemiller 1992; Winemiller and Rose 1992, 1993; Kopf et al. 2025). That is, in many ecosystems, species like buffalofishes and Freshwater Drum naturally occur in large numbers because long-lived adults are the lifeboat of the species and bridge environmental periods unfavorable for successful recruitment (Pereira 1992; Lackmann et al. 2021, 2023, 2024c). Large standing stock biomass in some of these taxa is exemplified by Swingle (1950), who reported results from full-scale rotenone surveys of eight large river sections in Alabama. These surveys were conducted before impoundment of most of these large rivers. Blue Catfish *Ictalurus fulcatus* and Channel Catfish (i.e., “rough fish escapees”) were the most abundant species in all but one river, and American Paddlefish *Polyodon spathula* (another “rough fish escapee”) was the dominant species in the other. In the Coosa River, Smallmouth Buffalo was the second most abundant species in terms of biomass. In the Tombigbee, Tensaw and Alabama Rivers, Freshwater Drum was the second most abundant species in terms of biomass. Therefore, catfishes, buffalofishes, drum, and paddlefish may naturally dominate biomass in some freshwater ecosystems. Paired with low replacement rates (P/B), this information suggests high harvest on these populations will diminish the large pools of standing stock, but this process may be cryptic, as has been demonstrated repeatedly in long-lived marine fishery stocks that were overfishes (Heppell et al. 2005). The overfishing of long-lived periodic strategist fishes (Winemiller and Rose 1992) may occur cryptically for years due to high initial biomass, but once declines are noticed directly by fisheries management the damage is often irreversible (Heppell et al. 2005). Similar dynamics of ‘hidden overharvest’ are observed in other fisheries, and in one example (Walleye), these patterns were detected explicitly because of a production-based approach (Embke et al. 2019). Another possibility in some situations is that harvest on these species is negligible, or recruitment occurs more regularly in some populations. However, longevity overfishing and age truncation of fish stocks is also problematic (Kopf et al. 2025; Slotte et al. 2025), and in long-lived sexually-size dimorphic species, unregulated or size-unrestricted harvest could negatively affect sex ratios (Heppell et al. 2005). Long-term fisheries community surveys with accurate age and sex data could be used to reveal the potential effects of high harvest limits on these species.

In this study, we deployed weighted GAMs to understand whether P/B predicted harvest limits in US states. We selected the GAM framework because it is agnostic to the shape of the P/B to harvest limit relationship. Furthermore, the P/B input data can be constrained to represent only the region of interest, making the model as realistic as possible for each scenario. Notably, across all states examined, relationships were consistently nonlinear. Though GAMs allowed flexible model shapes, the fits, while significant, remained weak (Table 1). We presented models for five US states, and the results reveal insights into the processes and culture of fish harvest limit setting in the USA. Despite their limitations, these relationships may still be useful for approximating initial bag limit estimates for native fishes so as to encourage further discussion. Using an example from Minnesota, we show how output from these models can be used to further align and check bag limits within the context of formal limit proposals. This ‘human-in-the-loop’ step is crucial for checking the accuracy of model outputs, especially given their marginal but significant fits. And ultimately, model outputs are only to be used as another tool or check within an otherwise larger process. In the Minnesota case, our model outputs happened to largely agree with proposed bag limits by Native Fish for Tomorrow (Winter 2024) (Table 2). These proposed bag limits were prepared by Native Fish for Tomorrow in anticipation of Minnesota’s Native Fish Bill and new native rough fish legislative category (Winter 2024). Board members proposed daily creel limits that considered perceived vulnerability to harvest referencing existing literature on life history, population trends, and ecosystem services, balanced with the yield of edible filets per individual. Even though these limits were developed independently, they are surprisingly close to our modeled limits. In this case, concordance between a science-based model and angler-derived limits provides a degree of confidence that proposed by Native Fish for Tomorrow make good biological sense and will result in improved resource management.

**Table 1.** Results of weighted GAM models for five US states and all states combined as presented in Figure 3. (Family: Gaussian, Method: REML).

| State | R<sup>2</sup><br>(adj) | Estimate<br>(SE) | edf | F-statistic | Deviance<br>explained | Parametric<br>coefficient<br>significance<br>(p-value) | Smoothing<br>term<br>significance<br>(p-value) |
| --- | --- | --- | --- | --- | --- | --- | --- |
| Georgia, GA | 0.50 | 28.72 | 1.08 | 10.47 | 53.90% | < 0.0001 | 0.0060 |
| Minnesota,<br>MN | -0.02 | 8.80 | 1 | 1.54 | 5.59% | 0.0021 | 0.2260 |
| Kentucky,<br>KY | 0.67 | 9.80 | 1.96 | 8.52 | 74.40% | 0.0002 | 0.0121 |
| New York,<br>NY | 0.181 | 13.53 | 1 | 4.29 | 23.4% | 0.0004 | 0.0574 |
| Oklahoma,<br>OK | -0.03 | 15.503 | 1 | 0.64 | 5.98% | 0.0048 | 0.4440 |
| All 5 States | 0.69 | 17.023 | 9.485 | 9.067 | 78.4% | < 0.0001 | < 0.0001 |

**Table 2.** Example of bag alignment exercise using Minnesota as a case study. Modeled bag limits from GAMs (see Methods) are compared to proposed limits from Native Fish for Tomorrow (NF4T), developed with no knowledge of the data or approaches in this study (see Discussion).

| Species or group | Average P/B (year<sup>-1</sup>) | Proposed limits from NF4T | Modeled bag limits |
| --- | --- | --- | --- |
| Mooneye and Goldeye, combined | 0.35 | 10 | 10 |
| Bowfin | 0.19 | 3 | 5 |
| Burbot | 0.31 | 4 | 8 |
| Bigmouth and Smallmouth buffalos, combined | 0.10 | 3 | 3 |
| Black Buffalo | 0.05 | 0 | 0 |
| White Sucker | 0.26 | 10 | 7 |
| Redhorses (Golden, Silver, Shorthead, River, Black, Greater), combined | 0.28 | 5 | 6 |
| Blue Sucker | 0.09 | 0 | 0 |
| Longnose Sucker | 0.13 | 3 | 4 |
| Carp suckers (Quillback, River, Highfin), combined | 0.68 | 5 | 9 |
| Freshwater Drum | 0.08 | 10 | 1 |
| Longnose and Shortnose gars, combined | 0.40 | 3 | 10 |
| Northern Hogsucker | 0.80 | 5 | 12 |

Our models also shed light on other biases occurring within our broader fisheries management systems. For example, ‘panfish’ species almost uniformly had bag limits that were poorly aligned with their P/Bs (bag limits too high). This misalignment is most clearly observed through the high residual values observed for these species in the model outputs. This discrepancy also supports a growing body of literature indicating that many panfish populations in the USA are overfished, at least with respect to size structure (Reed and Parsons 1999; Rypel 2015; Rypel et al. 2016; Clapp et al. 2021). In contrast, trout populations often exhibited more restrictive bag limits than predicted based on P/B (bag limits too low). In this case, the pattern is evidenced by extremely low/negative GAM residuals. Most trout populations have P/Bs of ∼1.0, indicating their biomass turns over roughly annually. Yet in most states, trout populations are managed using bag limits akin to species with P/B values of 0.2 (Walleye, Black Basses etc). Trout bag limits are therefore 80% lower than if you used the GAM to blindly set the limit, which may in turn help explain some of the overpopulation and density dependent issues observed in many US trout stocks (Beard Jr. 1999; Arismendi et al. 2024; Grossman et al. 2024; van Zyll de Jong 2025). The s-tip to our combined GAM shape is probably explained by two factors: 1) Harvest limits for Muskellunge set at one fish per day, even though most anglers will never harvest these fish, is essentially a trophy opportunity, but still a ‘safe’ policy because few will ever exercise it; 2) Harvest limits may still be too high for some game species, such as large Northern Pike *Exos lucius* and sturgeon as observed from their consistently high residual values in the GAMs. The history behind recreational fishing regulations in North America is interesting and often driven by sociological factors (Rypel et al. 2016). Above, we highlight some examples of misalignment in bag limits with P/B. There is much room for improvement in our fisheries management systems, and tools like those presented here can be used to probe into biases and harvest misalignment issues.

Our study, while an advancement in understanding and managing native fishees overall, represents a starting point. Ideally, management of any population is best when based on empirically-derived information for the population of interest rather than outputs from an empirical model. As seen in our relationship between P/B and maximum longevity, even though the fit is quite strong, there is still much variation. Therefore, use of these methods should not be viewed as a standalone replacement for the collection and study of the local ecology of these species and populations. As new information for data-limited species arises (e.g., Lackmann et al. 2021; 2022a; 2022b; 2023), use and reliance on these more oblique methods might be reduced. Secondly, in our study, we don’t address the potential of fishing contributing to size-structure problems (Grant et al. 2004; Diekert 2012; Rypel et al. 2016), which includes the nuance of how P/B also varies with size (Banse and Mosher 1980; Waters 1987; Randall 2002; Huryn and Benke 2007). There are major differences in P/B between stocks managed for size-structure versus those that are not; largely because of how removal of large fish truncates age-structure (Coble 1988; Hsieh et al. 2010), and thus probably lowers P/B (Banse and Mosher 1980). This issue is easily observed when assessing P/B of all Muskellunge in a population versus only harvestable size Muskellunge (often > 100 cm total length). Therefore, managing native fishes to prevent collapse is fundamentally different from managing these same stocks for quality and size-structure. As a result, regulations to ensure sustainability for a given native nongame species (what this study focuses on) might look quite different from regulations aimed at creating trophy angling opportunities for these same species. In the later example, limits would need to be reduced even lower as these actions would likely incorporate some form of length limits (Neumann and Allen 2007). The sustainability of these practices can still be assessed using a production-based approach; however, such goals must be built explicitly into the assessment. The study and management of Alligator Gar as a trophy angling opportunity largely supports this approach (Smith et al. 2018) and is somewhat akin to trophy Muskellunge management (Casselman 2007).

The long-term decline of native freshwater fishes worldwide indicates a stunning ineffectiveness of prevailing fisheries management systems (Xenopoulos et al. 2005; Walsh et al. 2011; Su et al. 2021; Vardakas et al. 2025). Although declining trends in North American biodiversity are less severe compared to other regions of the world, those which can be measured still indicate massive shifts, including risk of extinction for species in virtually all regions (Fagan et al. 2004; Freeman et al. 2005; Moyle et al. 2011). Freshwater fisheries management systems in the USA must change, and become more focused on protecting all species, not the select few historically favored by anglers. Standing in the way of improved management is a knowledge gap on the ecology of most species. In one survey of almost 200,000 scientific studies, Guy et al. (2021) found just 3% of articles focused on critically endangered fish; 82% of critically endangered species had no published articles. It is no wonder that we lack information needed to better manage our native fishes. The methods discussed in this paper provide one means of advancing management, despite limited data, for native fishes whose populations have been long-neglected or even antagonistically managed against. We encourage states to use or modify our approach, and to consider more protective limits on the harvest of native fishes. These changes will benefit fish populations, ecosystems, and human cultures for generations to come.

